# Cyborg Groups Enhance Face Recognition in Crowded Environments

**DOI:** 10.1101/357004

**Authors:** Davide Valeriani, Riccardo Poli

## Abstract

Recognizing a person in a crowded environment is a challenging, yet critical, visual-search task for both humans and machine-vision algorithms. This paper explores the possibility of combining a residual neural network (ResNet), brain-computer interfaces (BCIs) and human participants to create “cyborgs” that improve decision making. Human participants and a ResNet undertook the same face-recognition experiment. BCIs were used to decode the decision confidence of humans from their EEG signals. Different types of cyborg groups were created, including either only humans (with or without the BCI) or groups of humans and the ResNet. Cyborg groups decisions were obtained weighing individual decisions by confidence estimates. Results show that groups of cyborgs are significantly more accurate (up to 35%) than the ResNet, the average participant, and equally-sized groups of humans not assisted by technology. These results suggest that melding humans, BCI, and machine-vision technology could significantly improve decision-making in realistic scenarios.

## Introduction

Visual search is the process of looking for an item of interest in a scene. Everyone engages in visual search many times every day^1^. For example, when people look for their keys in a drawer, or when a bear hunts for fish in a river. Evolution has made animals very effective at searching in natural environments^2^. Yet, our performance is far from ideal^1,3^. For example, when the item of interest has a size that is inconsistent with the rest of the scene, we are more likely to miss it^4^. Phenomena such as inattentional blindness^5^ and illusory conjunctions^6^ lead us to miss objects or perceive them with wrong features.

These difficulties are present also when searching for a target face in pictures of crowded environments, a critical task in security and surveillance^7^. This challenging visual-search task requires humans to scan an image, detect faces, and compare them with the memory representation we have of a target person^8^. Even if our brain has dedicated regions to process face information^9^, humans still find this task demanding and difficult, even when highly-skilled^10^.

Searching for a target face in a crowded environment is a challenging task also for fully-automated computer-vision systems^11^, albeit not for the same reasons as humans. Firstly, a module of the system needs to scan the input image and extract all the faces with their location (*face detection*)^12^. Secondly, another module has to compare each extracted face against the image of the target face and return a degree of match (*face identification*)^12^. If the match is above a certain threshold, the image is classified as containing the target. Both face detection and face identification unavoidably produce false alarms and misses. Despite these difficulties, research has shown that deep neural networks do not suffer from some limitations of the human visual system^4^. In principle this means that computer-vision systems could become more accurate than the average human^13^, especially in tasks where visual information is presented for a brief amount of time. Unlike humans, the performance of computer-vision systems does not degrade over time (e.g., due to fatigue). However, it usually degrades way below that of humans when moving from constrained to realistic environments^14,15^.

To partially address the limitations of the human visual system, people can work together to search a scene. When making perceptual decisions, groups are generally more accurate than their members^16^. This “wisdom of crowds”^17^ has been found in a variety of tasks, ranging from estimating uncertain quantities^18^ to medical diagnosis^19^. However, there are circumstances in which groups decisions are *worse* than individual ones. For example, when the opinions of group members are too correlated^16^,when a strong leader dominates the discussion^20^, or when group interactions are not effective^21,22^.

The strategy used to aggregate individual opinions critically influences performance in group decision making. In the case of binary judgements, standard majority is the most straightforward strategy to use. However, in many circumstances it is suboptimal^23^. A well-chosen strategy could let groups make correct decisions even if the majority (but not all) of the members are wrong^24^. For instance, this could be achieved using a strategy consisting in weighing opinions on the basis of individual confidence^25^, which can be thought to approximate the probability of a decision being correct^26^. Nevertheless, an accurate assessment of the reliability of the decision is difficult to obtain. Humans may be miscalibrated in estimating their own confidence^27^, e.g., people may report high values of confidence when they made an incorrect decision. This miscalibration is even accentuated by experience^28^ and social influence^29^.

In principle, decision confidence could be more objectively assessed via means other than self-reporting. There is ample evidence that physiological correlates of decision confidence exist. For instance, several brain-activity patterns as recorded via electroencephalography (EEG) are known to provide information about decision confidence (e.g., the P300 event-related potential (ERP) and the error related negativity^30,31^), and visual search performance (e.g., the N2pc^32^ and N1^33^ ERPs). Moreover, it has been known for a long time that response times (RTs) negatively correlate with the confidence in a decision^34–36^. An important question is, however, whether these findings can be converted into practical systems to aid decision making.

In recent research, we developed a series of *hybrid Brain-Computer Interfaces* (hBCIs) to test whether it is indeed possible to decode the decision confidence of single users from their EEG signals, RTs and other physiological measurements. In all cases, the hBCI relies heavily on machine-learning technology, which is used to specialise the interface to each particular user and to learn to predict the probability that, in a certain trial, the user made a correct decision. The confidence estimates provided by the hBCI were then used to weigh individual opinions and obtain group decisions. That is, our hBCI did not make automated decisions, it only produced weights for integrating *behavioural* decisions provided by the operators. These hBCI-assisted group decisions were then compared to decisions obtained by other forms of opinion aggregation. After initial tests of our system with a simple visual-matching task^37^, which showed that hBCI-assisted groups were significantly more accurate than equally-sized groups based on standard majority, we progressed to much more realistic situations. For instance, we have shown that similar accuracy improvements could be obtained in visual search in crowded environments^38,39^, where participants were presented with cluttered scenes for 250 ms and had to decide whether a target object was present. The confidence estimated by our hBCI was also a more reliable predictor of correctness than the confidence reported by the participants themselves^39^. More recently, we have used our hBCI to assist humans searching for a target face in a crowded, indoor environment^40^, which is of critical importance in many domains and is the application area considered also in this article. Once again, even in this real-world situation, groups of humans assisted by our hBCI were significantly more accurate than traditional groups based on standard majority or reported confidence^40^. While this is encouraging, with or without the assistance of an hBCI, even trained operators find this task hard and very fatiguing.

Given advantages and limitations of both humans (alone or in a group, and with or without the assistance of an hBCI) and computer-vision systems, one may wonder whether we could usefully *combine* these together. At least in principle, within such “cyborgs”, automated algorithms could compensate for the weaknesses of humans and *vice versa*, thereby producing more reliable and accurate decisions. For example, in a situation where most users in a group missed a target face (e.g., because the target appeared in an unusual/unsuspected location), both traditional and hBCI-assisted groups would likely make a wrong decision. Conversely, a machine-vision algorithm might still identify the target with confidence. So, if it was included as a member in a group of humans, the machine-vision algorithm could potentially swing the group decision towards a correct decision.

This study explores the benefits and drawbacks of this idea. We used a state-of-the-art residual deep neural network (ResNet), our BCI estimating the decision confidence using only neural features, and human participants to create different types of cyborgs and investigate whether they can improve decision making and why. We evaluated these approaches in the context of the same face-recognition experiment as in^40^, where participants had to search an image of a crowded, indoor environment and decide whether or not it contained a target face.

## Methods

### Participants

Ten healthy participants (37.8 *±* 4.8 years old, seven females, all right-handed) with normal or corrected-to-normal vision and no reported history of epilepsy took part in the experiment after giving informed, written consents. Each participant was paid GBP 16 for taking part in the experiment plus an additional amount of up to GBP 4 proportional to the participant’s percentage of correct decisions. The latter was used to encourage participants to focus on the task and achieve maximum performance. This research received UK Ministry of Defence and University of Essex ethical approval in July 2014.

### Stimuli and procedure

The experiment consisted of six blocks of 48 trials. At the beginning of each block, participants were shown a cropped face image of the person to be considered as a target for that block and were asked to memorise it before starting the presentation of the trials in the block. Each trial (Figure 1) started with the presentation of a fixation cross for 1 s, followed by an image of a crowded scene presented for 300 ms in fullscreen, subtending approximately 14.4 degrees horizontally and 11.0 degrees vertically. Then, participants were presented with a display showing (again) the image of the target face and were asked to decide whether or not the target person was present in the scene, by clicking the left or the right mouse buttons, respectively, as quickly as possible. Finally, participants were asked to indicate, within 4 s, their subjective degree of confidence in that decision (0–100%) using the mouse wheel to vary the selected value in steps of 10%. Subjective confidence values were not used in this study as in preliminary analysis^40^ we found that these confidence estimates were less accurate and robust than the confidence estimated by the BCI from the brain signals.

**Figure 1.**
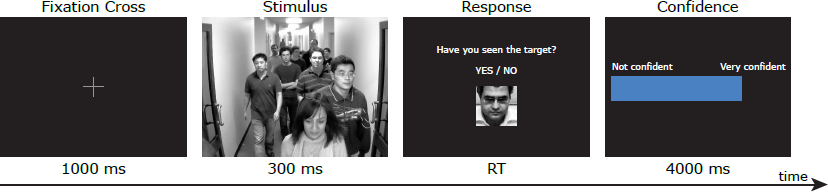
Sequence of stimuli presented in a trial.

The images used as stimuli were obtained from the sequences P2L S5 and P2E S5 of the ChokePoint video dataset^41^. The two sequences contained video streams of 700+ frames, representing 29 people (six females) walking indoor through two different portals. For each sequence, three videos were recorded from cameras positioned at the top-left (L), top-center (C), and top-right (R) of each portal, respectively. From each sequence, we selected one person as a “target”. Then, we randomly selected 48 frames out of the 700+ available, 12 where the “target” person was visible in the scene and 36 where it was not. Since each frame was represented by three images taken from the three viewpoints, we selected a total of 2 *×* 48 *×* 3 = 288 images. Each image, containing between 2 and 11 people, was converted to greyscale and had its histogram equalised.

In each block of the experiment, the images selected from a given sequence (1 or 2) and viewpoint (L, C or R) were presented. All six possible combinations of sequences and viewpoints have been tested. The order of presentation of the images in a particular block was the same for all participants. However, the order of presentation of the blocks was pseudo-randomly chosen for each participant, for counterbalancing reasons. In each block 25% of the images contained the target.

Participants were comfortably seated at about 80 cm from a LCD screen. Briefing, preparation and task familiarisation (via two blocks of 10 trials each) took about 45 minutes, while the experiment took roughly 25 minutes.

### Data recording and BCI confidence estimation

Neural data were recorded at 2048 Hz from 64 electrode locations with a BioSemi ActiveTwo EEG system. Each channel was referenced to the average of the electrodes placed on each earlobe and band-pass filtered between 0.15 and 40 Hz. Artefacts caused by ocular movements were removed with a standard correlation-based subtraction algorithm. From each trial, response-locked epochs starting 1 s before the user’s response and lasting 1.5 s were extracted from the EEG data, baseline corrected with the average voltage recorded in the 200 ms before the stimulus onset, and downsampled to 32 Hz. Each epoch was then associated to the class “correct” (confident) or “incorrect” (not confident) depending on whether the decision made by the participant in that trial was correct or not, respectively.

Common Spatial Pattern (CSP)^42^ filtering was used to transform the data in a subspace where the variance between the “correct” and “incorrect” classes was maximum. For each participant, we computed the CSP transformation matrix using the epochs in the training set, and used such a matrix to transform the data in the test set. The logarithm of the variances of the first and the last rows were computed and used as neural features.

For each participant *p*, we trained a logistic regression classifier to predict the decision confidence *w_p_*, i.e., the probability of a particular decision being correct. We used 8-fold cross-validation to ensure that the results were not unfairly affected by overfitting.

### ResNet architecture and training

We used a residual neural network (ResNet) (see `face_recognition` Python library available at https://github.com/ageitgey/face_recognition) to decide whether or not the target person was present in each image used as a stimulus in our experiment. The ResNet was formed of 29 convolutional layers (similarly to the ResNet-34 described by He et al.^43^), and had previously been trained on about 3 million faces derived from a number of datasets and tested on the “Labeled Faces in the Wild” benchmark^44^, where it achieved 99.38% accuracy.

In each trial of our experiment, the ResNet firstly scanned the input image to identify and extract individual faces using a pre-trained face-detection model integrated in the `face_recognition` library. The image of each face was then mapped to a 128-dimensional vector space where images of the same person are near to each other (i.e., face encoding). Then, the ResNetcomputed the difference between the encoding of each face and the encoding of the target face, normalised using the Frobenius norm. If any extracted face had a difference smaller than a threshold, *t*, the stimulus was labelled as “target”, otherwise it was labelled as “nontarget”. The threshold was set to the value that maximised the accuracy on the training set using 8-fold cross-validation (*t* = 0.535 *±* 0.016 across the folds).

In each trial, the confidence weight of the ResNet, *w_ResNet_*, was computed as follows:

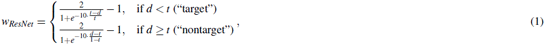

where *d* is the difference between the encodings of the face extracted from the image and the target face (normalised using the Frobenius norm). If more than one face was extracted from an image, only the face with the minimum *d* was considered. Figure 2 shows a plot of the function in Equation (1) for the average *t*. This formula ensures that the ResNet has substantially bigger confidence weights when the difference between the target face and the stimulus is very small or very big (confident decisions) than when it is close to the threshold (not confident decisions).

**Figure 2.**
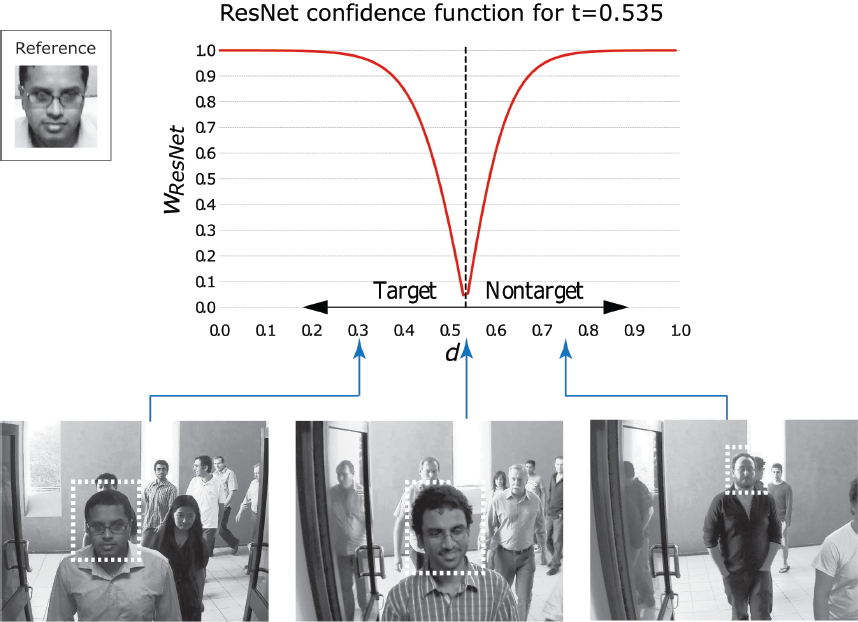
Function used with the ResNet to transform the difference between the encodings of the stimulus and target faces, *d*, into confidence weights for the average value of the threshold *t* over the eight folds. Examples of stimuli with *d*=0.301 (left), *d*=0.535 (middle), and *d*=0.750 (right) are also shown at the bottom. In each image, the bounding box of the face with the minimum *d* is highlighted. The reference image of the target person is shown on the top left.

Trials where the ResNet did not identify any face in the image were labelled as “nontarget”, given the prevalence of nontarget stimuli in the training set (i.e., 75%). In such trials, the corresponding weights *w_ResNet_* were set to 0, so that, in cyborg groups, the ResNet vote would be ignored and only the humans influenced the decision.

### Making group decisions

Groups of size *m* = 2*,…,*10 were formed off-line by combining the *N*=10 participants in all possible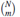are illustrated in Figure 3 and are described in detail below.

**Figure 3.**
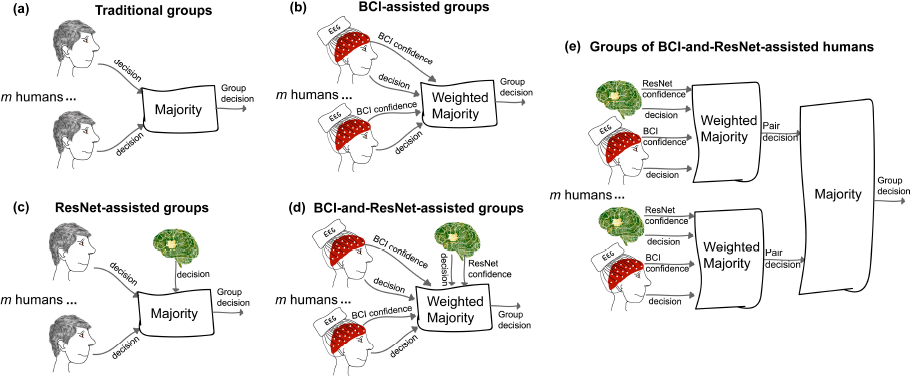
Summary of the different methods for obtaining decisions of groups of size *m*. (a) Traditional groups of human participants based on standard majority. (b) Groups of human participants weighing individual decisions with the confidence estimated by the BCI. (c) Traditional groups based on standard majority with an additional member being the ResNet. (d) BCI-assisted groups with the ResNet as an additional member, with its degree of confidence. (e) Groups where each participant is assisted by the BCI and paired with the ResNet, and the decisions of these pairs are then integrated using standard majority.

Traditional and BCI-assisted groups were composed only by human participants (Figure 3(a) and (b), respectively). In this case, the decision of a group of size *m* in a trial wasgiven by:

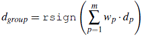

where *d_p_*= *−* 1 if user *p* decided the trial represented a “target” and *d_p_*= + 1 for a “nontarget”, *w_p_* is the corresponding weight, and rsign a randomising sign operator which returns +1 (“nontarget”) if its argument is positive, *−*1 (“target”) if it is negative, and randomly chooses between +1 and *−*1 if its argument is 0. Traditional group decisions were obtained using standard majority, i.e., *w_p_*=1 for all trials and users, while BCI-assisted group decisions used the decision confidence estimated by the BCI from the brain signals as weight *w_p_*.

ResNet-assisted and BCI-and-ResNet-assisted groups (Figure 3(c) and (d), respectively) were composed by *m* human participants plus the ResNet. Their decisions in a trial were given by:

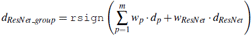

where *d_ResNet_* ∈ {−1,1} is the decision of the ResNet in the trial, and *w_ResNet_* is the corresponding weight. ResNet-assisted group decisions were obtained by using standard majority, i.e., *w_p_*=*w_ResNet_*=1 for all trials and participants, while BCI-and-ResNet-assisted group decisions were obtained by using the decision confidence estimated by the BCI from the brain signals as weight *w_p_*, and the confidence of the ResNet *w_ResNet_* as weight for its decision.

Finally, groups of *m* BCI-and-ResNet-assisted humans (Figure 3(e)) were obtained by firstly pairing each participant with the ResNet and computing their *d_ResNet_ group* decisions. These *m* decisions were then integrated using standard majority to obtain the decision of a group of BCI-and-ResNet-assisted humans:

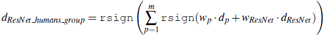

This last approach was suggested by literature for human groups^45^ showing that aggregating votes from smaller inner crowds outperforms pooling votes from larger groups of individual decision makers.

## Results

### The ResNet performs better than the average individual, but with lower sensitivity

The accuracy of the participants in the experiment was quite varied, ranging from 52.8% to 92.4%, with an average accuracy of 72.3% and a standard deviation (SD) of 12.0%. Their average specificity was 77.4% (SD=15.8%) and their average sensitivity was 56.9% (SD=10.9%). Hence, they recognised nontarget stimuli much better than target ones.

The ResNet had an average accuracy of 81.6%, a specificity of 96.8% and a sensitivity of 36.1%. Hence, it was substantially more accurate than the average participant. However, the ResNet was particularly good in assessing nontarget images (i.e., high specificity), but quite weak in detecting targets (i.e., low sensitivity) compared to the average participant. Since target stimuli appeared only in 25% of the trials, the ResNet could still achieve high accuracy, which was the metrics used to choose the threshold *t* (see *Methods*). In two nontarget trials and one target trial, the ResNet did not identify any face in the image and, therefore, classified the picture as “nontarget” (see *Methods*). Hence, only one error made by the ResNet was caused by its first step (i.e., extracting faces), which we could assume to be a trivial task for humans.

One may wonder whether a ResNet using a different metrics to optimise the choice of the threshold *t* could achieve better sensitivity. When choosing *t* in order to maximise the F1 score in cross-validation instead of the accuracy, the ResNet reduced its accuracy (80.9%) and specificity (92.1%), and boosted its sensitivity (47.2%). The average value of the threshold *t* over the eight folds was 0.572, while it was 0.535 when maximising accuracy. Similar results were obtained when maximising Cohen’s kappa to optimise *t* (average *t*=0.562, accuracy=80.6%, specificity=92.1%, and sensitivity=45.8%). In all cases, the sensitivity of the ResNet was worse than that of the average participant.

### The ResNet distinguishes itself from the crowd

To exhibit error-correction capability, groups require their members to make errors in different trials^46^. In fact, if all group members make the same, incorrect decision in a trial, the group will also make an incorrect decision. To assess error-correction capability of pairs, we used the Hamming loss, i.e., the fraction of trials in which the decisions of the two users were different. We computed the Hamming loss of each pair of participants, as well as the Hamming loss of each user when paired with the ResNet. We also computed the Hamming loss between the decisions of each user or the ResNet and the ideal user (i.e., one that is always correct), which corresponds to their error rates.

Participants had performance better than random and, therefore, their decisions were the same in the majority of the trials. However, the Hamming loss between two participants ranged from 12.5% and 47.2% (mean=30.67%, SD=7.92%). This suggests that the pairs exhibited error-correction capability in a large percentage of trials.

When looking at the Hamming loss between the ResNet and each participant, we found similar results. In this case, the Hamming loss ranged between 13.5% and 47.8% (mean=28.85%, SD=11.08%). A Wilcoxon signed-rank test showed no differences between the Hamming loss of humans and ResNet (*p*=0.508). Given its high performance, of course the ResNet had minimum Hamming loss with the participants with the highest accuracy (i.e., P5 and P6). Yet, even when paired with them, the pair still exhibited error-correction capability in more than 13% of the trials.

Different decision makers can be seen as nodes in a weighted graph, with the Hamming loss measure between pairs of decision makers representing the weight (dissimilarity) between corresponding nodes. Then, utilising a graph layout algorithm one can flattened the graph to obtain an intuition of similarities and differences in decision-making behaviour. Figure 4 shows the result of this process when using the program `neato` from the `Graphviz` library.

**Figure 4.**
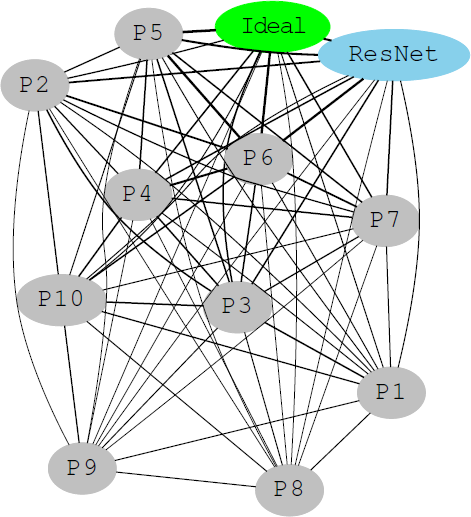
Graphical representation of the Hamming loss between the decisions of human participants (grey nodes), ResNet (light blue node) and ideal decision maker (green node). The weight of each edge is inversely proportional to the Hamming loss between the two nodes connected by that edge.

This diagram as well the previous analyses show how the ResNet behaves differently from virtually all other participants.

### Cyborgs have significantly higher performance that purely human groups

Figure 5 reports the performance of groups of increasing size using the five different methods for making group decisions analysed in this study (see Figure 3). The average performance of individuals and of the ResNet are also reported for reference.

**Figure 5.**
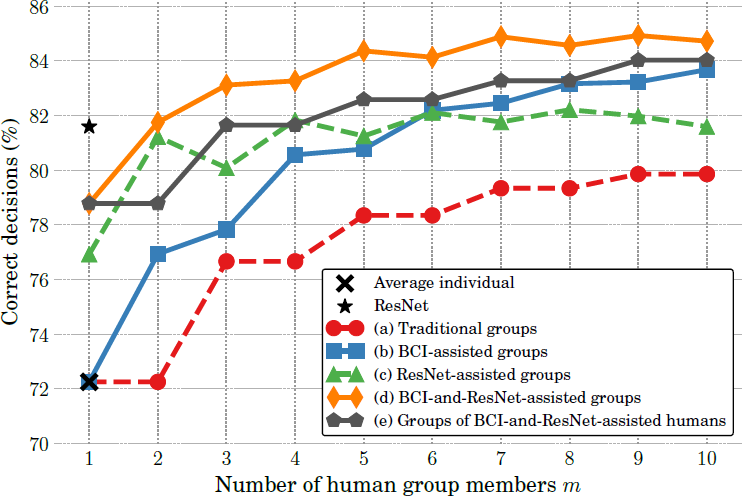
Average accuracy of individuals, ResNet, and groups of increasing size making decisions using the approaches described in Figure 3(a)–(e).

As one would expect^16^, the more opinions are integrated in the group decisions, the more accurate the group becomes. However, it is also apparent from the figure that the method chosen to integrate individual opinions and obtain group decisions is at least as important in determining group performance as the group size. In particular, it is clear that traditional groups of humans (red line in Figure 5) using standard majority are outperformed by all other four methods, regardless the group size.

Of course, one may expect methods which use some form of confidence estimation on a decision-by-decision basis to break ties, such as the BCI-assisted groups (blue line in Figure 5), to be superior to methods using simple majority and breaking ties randomly. However, the performance of ResNet-assisted groups (green line in Figure 5), which do not make use of confidence estimates, *appears* to contradict this expectation. Such groups are not only more accurate than equally-sized traditional groups, but also more accurate than BCI-assisted groups of sizes 2–5. This happens because the ResNet acts as an additional, very accurate member within the group (the ResNet is not only significantly better than the average participant, but only second to P5 and P6 in terms of performance). Since ResNet-assisted groups are based on standard majority, the smaller the group, the bigger the influence of the ResNet on the group decision. Indeed, the biggest improvement brought by ResNet-assisted groups over traditional groups occurs for *m*=2, where the addition of the ResNet allows us to turn most of the ties generated by the two humans into correct decisions. This results in a boost in accuracy of almost 9% over traditional pairs. For bigger group sizes, the difference in average performance between ResNet-assisted and traditional groups gradually diminishes. We should expect the performance of ResNet-assisted groups to eventually converge to that of traditional groups. Yet, even for the larger group sizes considered in our study, ResNet-assisted groups were still better that traditional groups by 2–3%.

Given the benefits provided by assisting groups with the BCI and the ResNet individually, it is not surprising to see that assisting groups with both (orange line in Figure 5) yields even better performance — in fact, the best performance over all other methods of obtaining group decisions.

Interestingly, when using groups of BCI-and-ResNet-assisted humans (grey line in Figure 5), groups were generally more accurate than ResNet-assisted groups (the only exception being groups of size *m*=2), but less accurate than BCI-and-ResNet-assisted groups. This indicates that, contrary to what is suggested in the literature for human groups^45^, aggregating votes from smaller inner crowds does not outperform pooling votes from larger groups of individual decision makers, when crowds are composed by humans and artificial agents.

To statistically validate the aforementioned observations, we used the Wilcoxon signed-rank test to perform pairwise comparisons of the performance of groups of increasing sizes using the five methods analysed in the study. Table 1 reports the two-sided *p*-values returned by the test. Statistical analyses for groups of size 10 was not possible as we only had one such group.

**Table 1.** Two-sided *p*-values of the Wilcoxon signed-rank test comparing the performance of: (a) traditional groups, (b) BCI-assisted groups, (c) ResNet-assisted groups, (d) BCI-and-ResNet-assisted groups, and (e) groups of BCI-and-ResNet-assisted humans. Values in bold represent *p<*0.05 confidence level.

| Group size | Sample size | (a) vs (b) | (a) vs (c) | (b) vs (c) | (c) vs (d) | (b) vs (d) | (d) vs (e) |
| --- | --- | --- | --- | --- | --- | --- | --- |
| 1 | 10 | 1.0 | $7.4 \times 10^{-2}$ | $7.4 \times 10^{-2}$ | $3.3 \times 10^{-1}$ | $1.5 \times 10^{-2}$ | 1.0 |
| 2 | 45 | **$1.1 \times 10^{-7}$** | **$5.5 \times 10^{-9}$** | **$7.9 \times 10^{-7}$** | **$3.6 \times 10^{-4}$** | **$1.2 \times 10^{-7}$** | **$1.1 \times 10^{-5}$** |
| 3 | 120 | **$6.0 \times 10^{-15}$** | **$8.7 \times 10^{-19}$** | **$5.9 \times 10^{-16}$** | **$1.1 \times 10^{-20}$** | **$1.4 \times 10^{-20}$** | **$2.9 \times 10^{-8}$** |
| 4 | 210 | **$1.5 \times 10^{-35}$** | **$3.3 \times 10^{-36}$** | **$4.8 \times 10^{-17}$** | **$1.1 \times 10^{-30}$** | **$7.8 \times 10^{-32}$** | **$5.0 \times 10^{-14}$** |
| 5 | 252 | **$2.4 \times 10^{-39}$** | **$7.9 \times 10^{-42}$** | **$5.8 \times 10^{-8}$** | **$7.1 \times 10^{-43}$** | **$5.1 \times 10^{-43}$** | **$6.9 \times 10^{-19}$** |
| 6 | 210 | **$3.3 \times 10^{-36}$** | **$3.2 \times 10^{-36}$** | $2.8 \times 10^{-1}$ | **$3.4 \times 10^{-33}$** | **$6.2 \times 10^{-33}$** | **$4.5 \times 10^{-19}$** |
| 7 | 120 | **$2.8 \times 10^{-21}$** | **$2.8 \times 10^{-21}$** | **$2.0 \times 10^{-10}$** | **$1.9 \times 10^{-21}$** | **$4.0 \times 10^{-21}$** | **$8.9 \times 10^{-13}$** |
| 8 | 45 | **$5.1 \times 10^{-9}$** | **$5.1 \times 10^{-9}$** | **$1.0 \times 10^{-6}$** | **$5.0 \times 10^{-9}$** | **$1.6 \times 10^{-8}$** | **$1.4 \times 10^{-5}$** |
| 9 | 10 | **$5.0 \times 10^{-3}$** | **$5.0 \times 10^{-3}$** | **$7.6 \times 10^{-3}$** | **$5.0 \times 10^{-3}$** | **$5.0 \times 10^{-3}$** | **$5.9 \times 10^{-2}$** |

As expected, BCI-assisted groups are significantly more accurate than equally-sized traditional groups based on standard majority for all group sizes – see “(a) vs (b)” column in Table 1. This confirms previous results obtained with this collaborative BCI both with the same set up (but using a combination of different neural features and reaction times)^40^, and in other visual-search experiments^39^. Moreover, ResNet-assisted groups were also significantly more accurate than traditional groups of only humans for all group sizes, as per “(a) vs (c)” column in Table 1. When comparing ResNet-assisted and BCI-assisted groups (“(b) vs (c)” column in Table 1), the test confirms that the former are significantly more accurate than the latter in small groups (i.e., *m*=2–5). Conversely, BCI-assisted groups are significantly better than ResNet-assisted groups for large group sizes (*m*=7–9). The two methods were on par for groups of size 6. For all group sizes *m*, BCI-and-ResNet-assisted groups (orange line in Figure 5) are significantly more accurate than groups assisted by only the ResNet or the BCI (“(c) vs (d)” and “(b) vs (d)” columns of Table 1, respectively). They are also significantly more accurate than groups of BCI-and-ResNet-assisted humans (“(d) vs (e)” column in Table 1).

### BCI and ResNet confidence correlates with correctness

Figure 6 shows the distribution of confidence values estimated by the BCI for the human participants (left) and the distribution of confidence values of the ResNet (right), for incorrect and correct decisions. The diagram on the left has been drawn with data from 799 incorrect trials and 2081 correct ones, while the diagram on the right has been obtained from 53 incorrect trials and 235 correct ones.

**Figure 6.**
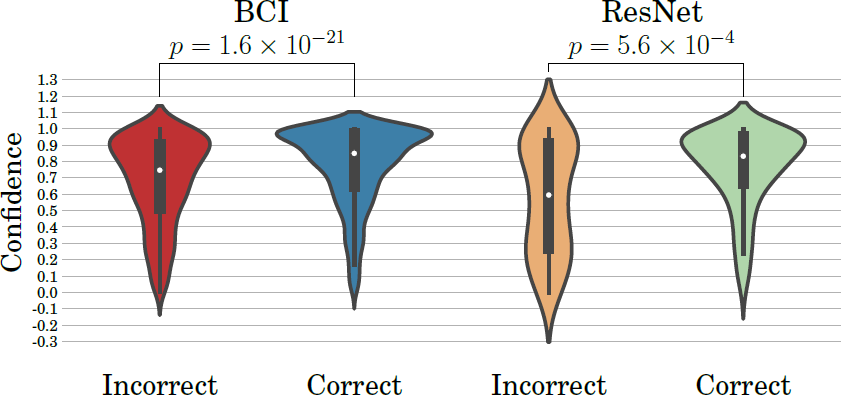
Distribution of confidence values for incorrect and correct decisions for the BCI (left) and the ResNet (right). The *p* values of a Kruskal-Wallis test comparing the correct and incorrect distributions are also reported.

Figure 6(left) indicates that, as we found in other experiments^39^, the BCI is able to assign markedly higher values of confidence to trials in which participants are correct than to trials in which they are incorrect. The median BCI confidence values for incorrect and correct trials are 0.747 and 0.850, respectively. The ResNet is also able to assess its degree of confidence via Equation 1 (see *Methods*) and its confidence values are markedly lower in incorrect trials (median=0.595) than in correct ones (median=0.833). These differences are significant as indicated by the *p* values (reported in the figure) of a Kruskal-Wallis test comparing the incorrect and the correct distributions for each type of confidence.

We should note that the differences between the median values of correct and incorrect trials are much bigger for the ResNet (0.238) than for the BCI (0.103). From Figure 6 it is clear that, for both the BCI and the ResNet, incorrect trials have a much sparser distribution than correct ones. Indeed, the standard deviations of the BCI confidence for incorrect and correct trials are 0.268 and 0.239, respectively, while for the ResNet they are 0.334 and 0.241, respectively, and these differences are significant (Levene test *p*=2.9 *×* 10*^−^*^7^ and *p*=9.3 *×* 10*^−^*^6^, respectively). Once again, the difference in standard deviations between correct and incorrect confidence values is bigger for the ResNet.

Taken together, these statistical differences suggest that not only is the ResNet more correct than the average human participant, but it is also better in estimating its confidence than the BCI is in estimating the confidence of the average human participant.

Of course, this is not entirely unexpected. The BCI bases its confidence estimates on the brain signals of the users, hence facing two major challenges that affect its certainty. Firstly, EEG signals are generally affected by a high level of noise caused by various sources (e.g., muscular activity, electrical interference, etc.). Secondly, humans participants may be overconfident or underconfident, two behaviours which might in turn also impact their brain patterns, thereby making it potentially difficult for our algorithms to reliably infer the probability of a decision being correct. While also the ResNet decisions and confidence are affected by noise from different sources (e.g., partial occlusion of the target face), the fact that it is based on computer-vision algorithms and Equation (1) makes the ResNet more reliable.

Irrespective of the reasons for the differences in confidence distributions between ResNet and BCI-assisted humans reported in Figure 6, due to such differences, in a scenario where a BCI-assisted human works in pair with the ResNet and there is a tie, the human is likely to break the tie in his/her favour. While this may or may not be optimal from a group decision perspective, it may make working like a cyborg much more acceptable to the human in the team.

## Discussion

A cyborg is typically defined as the combination of a human and a machine, which is either invasively embedded inside the human or replaces/extends a body part, with the aim of restoring some function or augmenting human capabilities. These types of cyborgs were once only the domain of science fiction, but are nowadays becoming a reality thanks to the advancements in neural engineering and prosthetics. However, we believe that the terms cyborg and cyborg group are also appropriate to describe non-invasive combinations of one or more humans and one or more machines, such as the systems described in this paper to augment performance in face recognition.

Of course, such systems can also be considered as forms of Human-Machine Interaction (HMI), consisting of a machine analysing information, and humans using this analysis to make informed decisions. In this area fall hybrid BCIs^47^ that allow users to control certain parts of an external device with their mind, while other parts are controlled by another system. In this scenario, we generally refer to *shared-control* systems, where the artificial component of the HMI generally takes care of all low-level decisions^48^. For example, when driving a car^49^ or controlling a wheelchair^50^ with a hybrid BCI, the autonomous control system usually performs the obstacle avoidance, while the BCI, relying on noisy brain measures, only issues high-level commands^51^. However, in those scenarios humans and machines do not fully interact: they simply control different aspects of the device.

A more recent and promising avenue of research has been looking at Human-Machine Cooperation (HMC), which further extend HMI to allow the human and the machine to cooperate on a more equal footing and at a higher (cognitive) level^52^. This novel approach could allow us to create a team of human and artificial operators with no a priori dichotomy between high-level (human) and low-level (machine) elements. Instead, the contribution of each operator to the control strategy or the decision-making process depends on a time-dependent and situation-dependent assessment of each actor’s strengths and weaknesses. It is clear that in tasks which are challenging for both humans and machines, joining forces may allow us to significantly improve performance. However, it is not clear how the cooperation can be made to work practically.

The cyborgs and cyborg groups proposed in this study represent operational solutions to the problem of how HMC could be implemented in an uncentralised and self-organising manner in the domain of decision making. The simplest solution would be creating groups where humans and machines have an equal weight in the decision making (majority-based groups). In our face-recognition task, the ResNet achieved significantly better performance than the average participant, hence allowing ResNet-assisted groups to significantly improve performance over traditional groups of humans. However, this approach is situation-independent, i.e., it is not able to discriminate in which cases the machine or the human should be trusted more.

Cyborg group performance could be significantly boosted by assessing the decision confidence of team members (computers or humans) in each decision and weighing their decisions accordingly. We have shown that machine-vision algorithms could be designed that also estimate their degree of confidence, while BCIs could be used to decode the decision confidence of humans from their brain activity. These BCI-and-ResNet-assisted groups fully implement the HMC strategy, allowing to decide the contribution to the group decision of each team member (human or machine) on a decision-by-decision basis, depending on the degree of confidence of each actor. Moreover, contrary to what has been suggested in the literature for human groups^45^, cyborg groups function better when human and machine decisions are integrated at the same level, instead of integrating decisions of human-machine pairs.

The cyborgs proposed in this study also takes into account ethical and liability issues concerning automated decision making. We have shown that the confidence values provided by the BCI for humans are higher than the confidence of the ResNet, leading to humans counting often more than the ResNet in uncertain decisions. When cyborg groups are used at an operational level, it is ethically simpler (albeit not necessarily optimal) to let humans decide in cases where there is no clear majority of opinions between humans and ResNet. However, when humans in a group cannot agree on a decision, it is more reasonable to trust fully-automated systems than flipping a coin to break ties.

Overall, all strategies explored in this study on how to integrate humans and machines have shown significant advantages in face-recognition performance over traditional groups of only humans. When fully-automated machine-vision algorithms are not reliable enough, BCIs could be used to significantly boost the performance of groups of human operators. In other situations, pairing machines and BCI-assisted humans may provide the best performance. Future work will verify these findings with other tasks where automated algorithms could be developed.

## Acknowledgements

The Authors acknowledge support of the UK Defence Science and Technology Laboratory (Dstl) and Engineering and Physical Research Council (EPSRC) under grant EP/P009204/1. This is part of the collaboration between US DOD, UK MOD and UK EPSRC under the Multidisciplinary University Research Initiative. The Authors also acknowledge support by the Defence and Security PhD programme through Dstl.

## Author contributions statement

D.V. and R.P. conceived and designed the study. D.V. collected and analysed the data. D.V. and R.P. interpreted the data and wrote the manuscript.

## Additional information

Competing financial interests: The authors declare no competing financial interests.

